# Functional expression of the ATP-gated P2X7 receptor in human iPSC-derived neurons and astrocytes

**DOI:** 10.1101/2021.03.28.437391

**Authors:** Jaideep Kesavan, Orla Watters, Klaus Dinkel, Michael Hamacher, Jochen H.M. Prehn, David C. Henshall, Tobias Engel

**Affiliations:** Department of Physiology & Medical Physics, Royal College of Surgeons in Ireland, University of Medicine and Health, Dublin D02 YN77, Ireland; Lead Discovery Center GmbH, Otto-Hahn-Straße 15, 44227 Dortmund, Germany; Affectis Pharmaceuticals AG, Otto-Hahn-Straße 15, 44227 Dortmund, Germany; FutureNeuro, Science Foundation Ireland Research Centre for Chronic and Rare Neurological Diseases, RCSI, Dublin D02 YN77, Ireland

**Keywords:** P2X7 receptor, human iPSC-derived neurons and glia, BzATP-evoked Ca^2+^ fluctuations

## Abstract

The P2X7 receptor (P2X7R) is a cation membrane channel activated by extracellular adenosine 5′-triphosphate. Activation of this receptor results in numerous downstream events including the modulation of neurotransmission, release of pro-inflammatory mediators, cell proliferation or cell death. While the expression of P2X7Rs is well documented on microglia and oligodendrocytes, the presence of functional P2X7Rs on neurons and astrocytes remains debated. Furthermore, to date, functional studies on the P2X7R are mostly limited to studies in cells from rodents and immortalised cell lines expressing human P2X7Rs. To assess the functional expression of P2X7Rs in human neurons and astrocytes, we differentiated human-induced pluripotent stem cells (hiPSCs) into forebrain cortical neurons that co-express FOXG1 and βIII-tubulin as well as S100 β-expressing astrocytes. Immunostaining revealed prominent punctate P2X7R staining on the soma and processes of hiPSC-derived neurons and astrocytes. In addition, our data show that stimulation with the potent nonselective P2X7R agonist BzATP induces robust calcium rises in hiPSC-derived neurons and astrocytes, which were blocked by the selective P2X7R antagonist AFC-5128. Together, our findings provide evidence for the functional expression of P2X7Rs in hiPSC-derived forebrain cortical neurons and astrocytes demonstrating that these cells offer the potential for investigating P2X7R-mediated pathophysiology and drug screening *in vitro*.

## 1. Introduction

The ATP-gated purinergic P2X7 receptor (P2X7R) plays a pivotal role in ATP-mediated signal transmission in the brain. ATP may be released from nerve terminals or glial cells by exocytosis [1] or from astroglial cells via non-exocytotic mechanisms. ATP has also been shown to leak through the damaged neuronal or glial plasma membrane during brain injury [2, 3]. When ATP binds to the extracellular domain of the P2X7R, the channel opens allowing the permeation of small cations such as Na^+^, Ca^2+^, and K^+^ [4]. In addition, P2X7Rs are also believed to form a non-selective “macro pore” which allows the influx of large-molecular-weight molecules [4]. Activation of P2X7R evokes glutamate release from presynaptic nerve terminals and from astrocytes leading to excitotoxic effects [5], while activation of P2X7R on astrocytes and microglia has been shown to lead to the release of Interleukin-1β (IL-1β), IL-6, and Tumor necrosis factor-α (TNF-α), triggering neuroinflammation [6]. The P2X7R has attracted considerable interest as a potential target for various central nervous system (CNS) disorders including epilepsy, and neurodegenerative diseases such as Parkinson’s disease, Alzheimer’s disease and multiple sclerosis as well as non-CNS diseases such as migraine and neuropathic pain [7-9]. P2X7R protein expression has been found to be highly up-regulated in the brain of patients with neurological diseases such as Huntington’s disease [10], intractable temporal lobe epilepsy (TLE) [11] and in cortical lesions of patients with focal cortical dysplasia [12].

Despite the 80% sequence homology between human and murine P2X7Rs, differences in the receptor sensitivity towards various ligands have been described. For example, the human P2X7R has been shown to be 10–100 times more sensitive to the non-selective agonist 2′,3′-O-(benzoyl-4-benzoyl)-adenosine 5′-triphosphate (BzATP) compared to the murine ortholog [13-15]. KN-62 (1- [N,O-bis(5-isoquinolinesulphonyl)-N-methyl-L-tyrosyl]-4-phenylpiperazine), a potent antagonist for human P2X7Rs has been shown to be inefficacious at rat P2X7Rs [16]. Likewise, human P2X7Rs subjected to ivermectin underwent positive allosteric modulation while modest results were seen at rodent P2XRs, suggesting a species-specific mode of action [17]. Thus, P2X7Rs exhibit remarkable species-specific differences in the pharmacological properties including agonist and antagonist potency and sensitisation which could account for the failure in clinical translation of the results from animal models [13]. Consequently, there is an increasing need for the investigation of the expression and function of P2X7Rs in human brain-relevant models. Resected human primary brain tissue is very scarce and often shows alterations associated with pathology. Advances in human induced pluripotent stem cell (hiPSC) technology has led to the differentiation of region-specific human neurons, astrocytes, oligodendrocytes and microglia from somatic cells [18-21]. These iPSC-derived human neurons and glia have demonstrated tremendous potential for the investigation of human neuronal development *in vitro*, pharmacological screening and assessing phenotypic alterations in patient-specific iPSC-derived neural cells [22-27].

To date, only a few studies have addressed the functional expression of the P2X7R in human neurons and astrocytes compared with work on rodent counterparts [28-30]. The main purpose of this study was to characterize the functional expression of P2X7Rs in hiPSC-derived neurons and astrocytes. Here, we demonstrate the expression of P2X7Rs using immunocytochemical analysis and BzATP-evoked Ca^2+^ transients in hiPSC-derived neurons and astrocytes.

## 2. Materials and Methods

### Culture and Differentiation of hiPSCs

hiPSCs HPSI0114i-eipl_1 (ECACC 77650081; Culture Collections, Public Health England, UK) at passage 32 were maintained under feeder-free conditions on vitronectin (STEMCELL Technologies, British Columbia, Canada)-coated 6-well plates in E8 medium (Thermo Fisher Scientific, Massachusetts, USA). hiPSCs were dissociated by using 0.5 mM EDTA for 2 min at 37°C, and reseeded at the density of 1 × 10^4^ cells per cm^2^. Cortical differentiation of hiPSCs was carried out as described previously, with minor modifications [31]. For the neural induction of hiPSCs, approximately 24 h after splitting, culture medium was switched to Gibco PSC Neural Induction Medium (Thermo Fisher Scientific, Massachusetts, U.S.A.) containing Neurobasal medium and Gibco PSC neural induction supplement. Neural induction medium was changed every other day from day 0 to day 4 of neural induction and every day thereafter. At day 10 of neural induction, primitive NSCs (pNSCs) were dissociated with Accutase (Thermo Fisher Scientific, Massachusetts, USA) and plated on Geltrex-coated dishes at a density of 1 × 10^5^ cells per cm^2^ in NSC expansion medium containing 50% Neurobasal medium, 50% Advanced DMEM/F12, and 1% neural induction supplement (Thermo Fisher Scientific, Massachusetts, U.S.A.). pNSCs were passaged on the 4^th^ day at a 1:3 split ratio to derive NSCs and cells at passage 4 were used for the differentiation of neurons and glia. For the differentiation of neurons, NSCs were plated on Geltrex-coated coverslips at a density of 5×10^4^ cells per cm^2^ in neuronal differentiation medium consisting of neurobasal medium and DMEM-F12 (1:1), with 2% B-27 supplement, 1% N2 supplement, 1% L-glutamine, 1% nonessential animal acids, 20 ng/ml brain-derived neurotrophic factor (BDNF), 20 ng/ml glial cell-derived neurotrophic factor (GDNF) (all from Thermo Fisher Scientific, Massachusetts, USA), 100 ng/mL cAMP, 100 µM L-ascorbic acid and penicillin/streptomycin (all from Merck, Missouri, United States). The culture medium was changed every 2–3 days. For astrocyte differentiation, pNSCs were plated onto Geltrex-coated coverslips at a density of 5 × 10^4^ cells per cm^2^ in an astrocyte differentiation medium (DMEM supplemented with 1% fetal bovine serum, 1% sodium pyruvate, 1% non-essential amino acids, 0.5% G-5 supplement (all from Thermo Fisher Scientific, Massachusetts, USA)) for 5 days. The astrocytes were passaged at a split ratio of 1:2 and medium was changed every 3 days. Astrocytes at passage 7 to 9 were used for experiments.

### Immunocytochemistry

NSCs, neurons and astrocytes cultured on coverslips were fixed with 4% paraformaldehyde for 15 min. After 3 washes in phosphate buffered saline, cells were permeabilized with 0.1% Triton for 20 min and blocked in 1% BSA for 30 min. Cells were incubated at 4°C overnight in primary antibody diluted in 1% BSA. The cells were then incubated in secondary antibody diluted in 1% BSA for 1 h at room temperature. The following primary antibodies were used: mouse anti-SOX2 (R&D Systems, Minneapolis, MN, USA), rabbit anti-nestin, mouse anti-PAX6, rabbit anti-FOXG1 (all from Abcam, Cambridge, UK), mouse anti-β-tubulin III (Biolegend, San Diego, CA) and rabbit anti-P2X7R (Alomone Labs, Jerusalem, Israel). The respective secondary antibodies were conjugated to Alexa Fluor 488 or Alexa Fluor 594 (Thermo Fisher Scientific, Massachusetts, USA). Coverslips were mounted and images were acquired using a Leica DM4 B fluorescence microscope.

### Patch-clamp electrophysiology

NSC-derived neurons differentiated for 2–8 weeks *in vitro* were selected for patch-clamp experiments. Whole cell voltage-clamp recording was carried out with a Multiclamp 700 B amplifier (Molecular Devices, California, USA) which was interfaced by an A/D-converter (Digidata 1550B, Molecular Devices, California, USA) to a PC running pClamp software (Version 11, Molecular Devices, California, USA). The signals were low-pass filtered at 2 kHz or 10 kHz and sampled at 10 kHz or 50 kHz. Pipette electrodes (G150T-4, Hardward Apparatus, Massachusetts, USA) were fabricated using a vertical puller (Narishige PC-100, Tokyo, Japan). All recordings were performed at 32°C in a bath solution containing (in mM): 135 NaCl, 3 KCl, 2 CaCl2, 1 MgCl2, 10 Hepes and 10 glucose (pH 7.2; osmolality 290–300 mmol/kg). For recordings of currents, the patch pipette contained the following solution (in mM): 120 potassium gluconate, 20 KCl, 10 NaCl, 10 EGTA, 1 CaCl2, 4 Mg ATP, and 0.4 Na GTP, and 10 HEPES (pH 7.2, osmolality 280–290 mmol/kg). Spontaneous action potential firing was detected in loose-patch configuration with patch pipettes filled with the bath solution.

### Drug application

Stock solutions of BzATP (Alomone Labs, Jerusalem, Israel) and AFC-5128 (Affectis Pharmaceuticals AG, Dortmund, Germany) were diluted and applied in HEPES-buffered extracellular solution. The drug solutions were delivered to the recorded cells by a valve-controlled fast multibarrel superfusion system with a common outlet approximately 350 µm in diameter (Automate Scientific, California, USA). The application tip was routinely positioned approximately 1 mm away from and ∼ 50 µm above the surface of the recorded cells. A computer connected to Digidata 1550B controlled the onset and duration of each drug application. The time course of the drug application was calibrated using fluorescein dye.

### Calcium Imaging

Cells were loaded at 37°C with fluo-4 (Abcam, Cambridge, UK) or Cal-520 (AAT Bioquest, California, USA) by incubation with the acetoxymethyl (AM) ester form of the dye at a final concentration of 2 μM in culture media. After 45 min, cells were washed several times with dye-free HEPES-buffered saline solution and transferred to an imaging chamber on a microscope (Zeiss Axio Examiner, Jena, Germany) equipped with a Zeiss 40x water immersion objective. Zen Blue imaging software (Carl Zeiss, Jena, Germany) was used for hardware control and image acquisition, and image analysis was performed using ImageJ (NIH, Maryland, USA). All imaging experiments were performed at 32°C in a low divalent cation containing bath solution with the composition (in mM): 135 NaCl, 3 KCl, 0.1 CaCl_2_, 0 MgCl_2_, 10 HEPES and 10 glucose (pH 7.2; osmolality 290–300 mmol/kg). Images were acquired at 4 Hz. Background fluorescence was measured from the cell-free area outside the soma of interest in each frame of every time series. Region of interests (ROIs) were manually drawn around the soma and baseline fluorescence intensity (F0) was determined by averaging 14 frames preceding the cell’s exposure to BzATP, and the time course of normalized fractional dye fluorescence [ΔF/F0] was obtained, where ΔF equals F(t)-F0.

### Data analysis

The data are expressed as mean ± SEM. Statistical analysis was performed by Student’s t-test. P < 0.05 was considered statistically significant. All statistical analysis was performed with GraphPad Prism 9 software (GraphPad Software, San Diego, CA, USA).

## 3. Results

### 3.1. Differentiation of forebrain cortical neurons and astrocytes from hiPSC-derived primitive neural stem cells

In order to assess the functional expression of P2X7Rs in a human brain-relevant cellular model, we differentiated cortical neurons and astrocytes from hiPSCs that were maintained on vitronectin-coated plates in E8 medium for 2 passages. The hiPSCs were passaged with 0.5 mM EDTA at a split ratio of 1:5. Cells maintained under these conditions were characterized by undifferentiated morphology with round and clear edges (Figure 1A).

**Figure 1.**
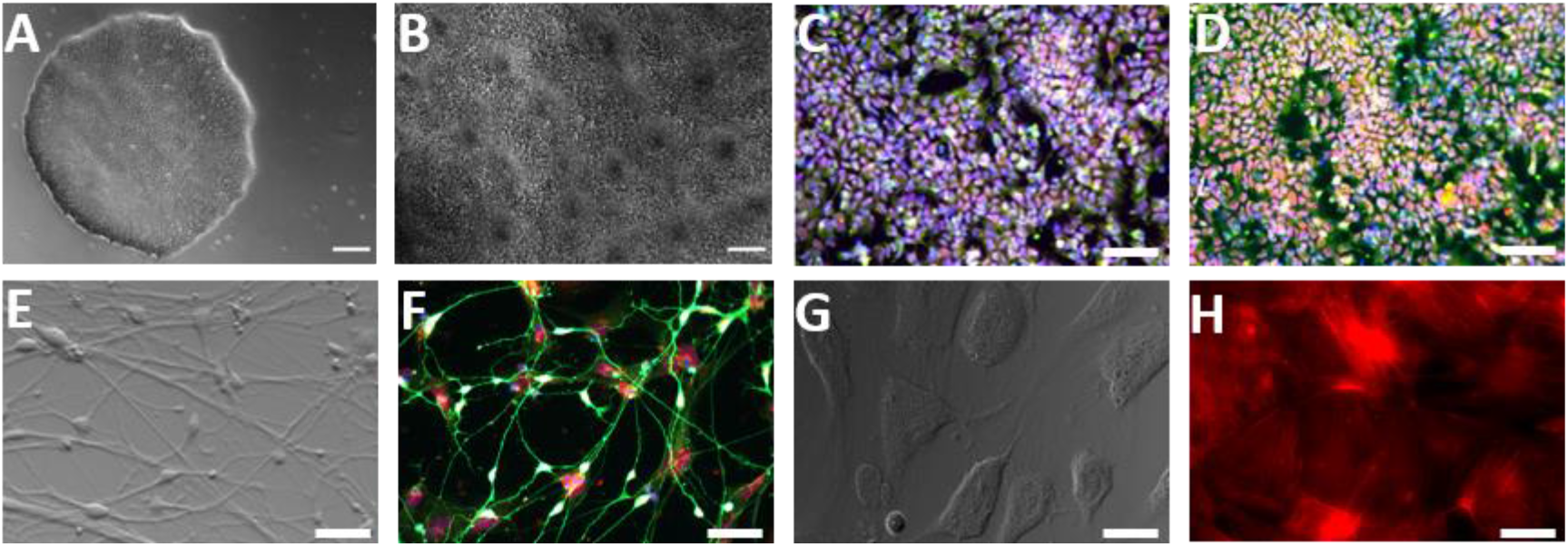
Differentiation of forebrain neurons from hiPSCs. (**A**) Brightfield image of hiPSC colony cultured in feeder-free conditions at day 3 *in vitro*. (**B**) pNSCs derived from hiPSCs at day 7. (**C**) Expression of neural stem/progenitor markers nestin (green) and SOX2 (red) in NSCs. Cell nuclei were stained with Hoechst (blue). (**D**) The NSCs co-express neural stem/progenitor markers nestin (green) and dorsal telencephalic progenitor marker PAX6 (red). Cell nuclei were stained with Hoechst (blue). (**E**) NSC-derived neurons at two weeks of differentiation *in vitro*. (**F**) Differentiated cells were stained with antibodies against the forebrain marker FOXG1 (red) and the neuronal marker, β-III tubulin (green). Cell nuclei were stained with Hoechst (blue). (**G**) Brightfield image of differentiated pNSCs in astrocyte induction media. (**H**) Differentiated astrocytes express astrocyte marker S100β. Scale bars 100 µm (A, B, C and D) or 50 µm (E, F, G and H).

Next, we differentiated hiPSCs to primitive neural stem cells by dissociating the hiPSC colonies using accutase and plating onto Geltrex-coated dishes in neural induction medium for 10 days resulting in a population of pre-NSCs (Figure 1B). The medium was changed every other day for the first 4 days and every day thereafter. Once confluent, cells were passaged 1:3 and could be cryopreserved in liquid nitrogen. At passage 4, the cells exhibited a homogeneous morphology and immunocytochemical analysis revealed the expression of specific markers of mature NSCs such as Nestin, SOX2 (Figure 1C) and the dorsal telencephalic progenitor marker PAX6 (Figure 1D). For neuronal differentiation, NSCs at passage 4 were plated on Geltrex-coated plates in neuronal differentiation medium. At 14 days *in vitro*, cells exhibited neuronal morphology and were positive for the neuron-specific cytoskeletal marker β-III-tubulin and forebrain marker FOXG1 (Figure 1E-F). For the differentiation of astrocytes, dissociated NSCs were plated onto Geltrex-coated plates at a density of 5 × 10^4^ cells per cm^2^ in an astrocyte differentiation medium and confluent cultures were passaged every 4-5 days at a ratio of 1:2, and medium was changed every 2–3 days. Immunocytochemical analysis at passage 7 confirmed the expression of the astrocyte marker S100β in cells maintained in astrocyte differentiation mediaum (Figure 1 G-H).

Thus, our data demonstrate efficient induction of pNSCs with the potential for differentiation into forebrain cortical neurons and astrocytes.

### 3.2. In vitro differentiated hiPSC-derived cortical neurons exhibit fundamental electrophysiological properties

For the functional assessment of the differentiated neurons, NSCs were plated on a monolayer of hiPSC-derived astrocytes from the same hiPSC line and maintained for 2 to 8 weeks. Using whole-cell patch-clamp recordings from individual neurons, we first confirmed that NSC-derived cortical neurons differentiate as functional neurons exhibiting fundamental electrophysiological properties. The recordings demonstrated the presence of fast activating and inactivating inward currents and slow and sustained outward currents resembling voltage-gated Na^+^ (Na_v_) and K^+^ (K_v_) channels (Figure 2 A-B).

**Figure 2.**
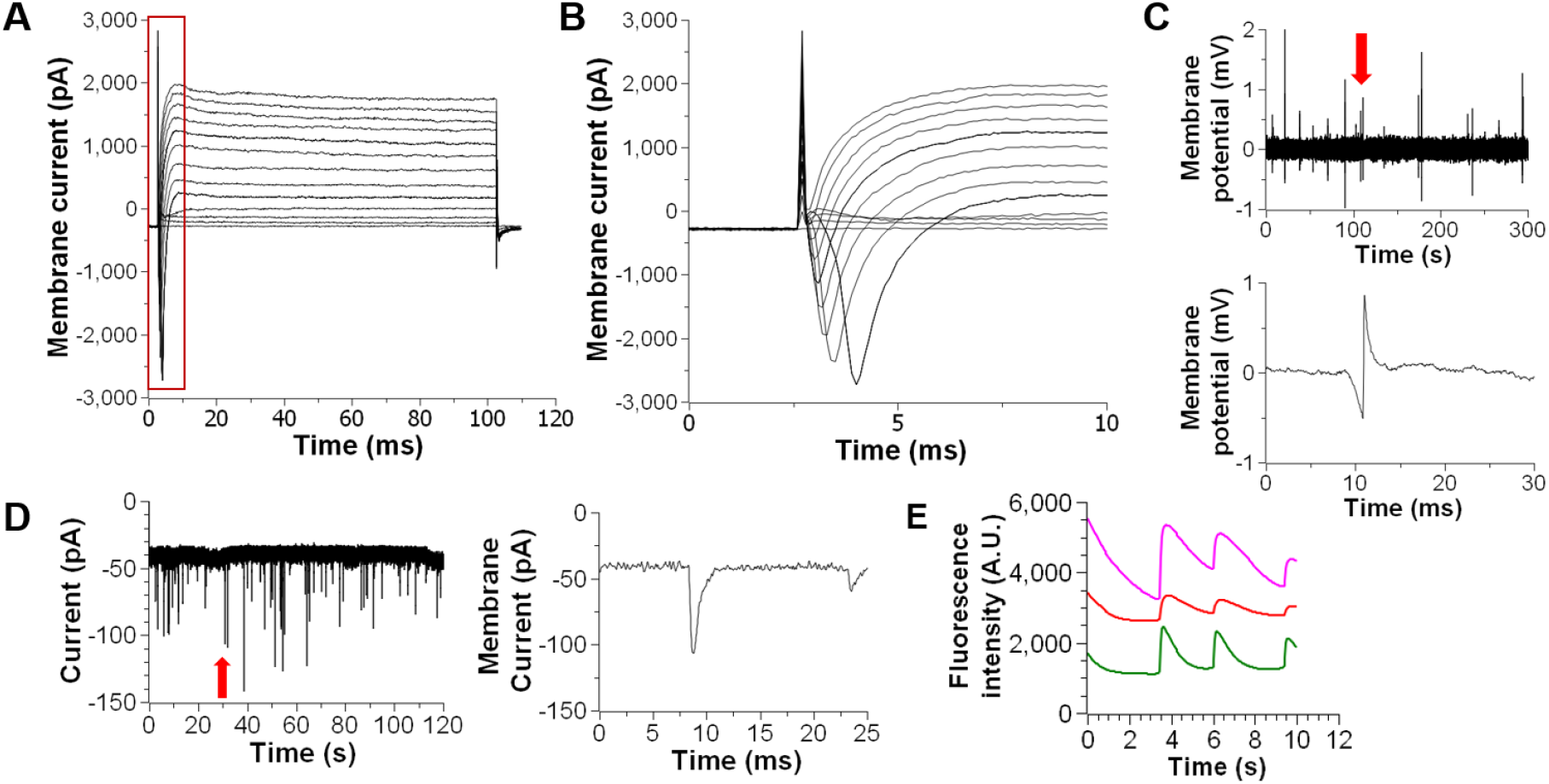
Fundamental electrophysiological properties of hiPSC-derived neurons. (**A**) Representative traces of fast-inactivating inward currents and sustained outward currents recorded in voltage-clamp mode. (**B**) Expanded view of the boxed region of inward and outward currents. (**C**) Action potentials recorded in loose-patch configuration. One action potential indicated by the red arrow is shown at expanded scale at the bottom. (**D**) Spontaneous postsynaptic currents (sPSCs) recorded in whole-cell configuration from hiPSC-derived neural network. Expanded view of the sPSC (right). (**E**) Example of spontaneous synchronized [Ca^2+^]_i_ transients in Fluo-4 loaded neurons.

Demonstrating neuronal maturity, non-permeating loose-patch recordings confirmed spontaneous action potential firing of neurons (Figure 2C). Another hallmark of a mature neuronal network is the formation of functional synapses. Whole-cell voltage clamp recordings revealed spontaneous postsynaptic currents (sPSCs) at 21-28 days *in vitro* differentiation (Figure 2D). In addition, synchronous Ca^2+^ transients measured by oscillations in Fluo-4 fluorescence were also evident in differentiated neurons indicating a synaptically connected network (Figure 2E).

Therefore, pNSCs terminally differentiate to neurons *in vitro* and exhibit fundamental electrophysiological properties including action potential firing and synaptic transmission.

### 3.3. Expression of P2X7Rs in hiPSC-derived neurons and astrocytes

To investigate if P2X7Rs are expressed in hiPSC-derived neurons, immunocytochemistry was performed at 8 h after plating the NSCs. At this stage, the differentiated cells with neuronal morphology expressed the neuronal marker β-III tubulin (Figure 3 A,B). In double-labelling experiments using the neuronal marker β-III tubulin and P2X7R-specific affinity purified antibody, expression of P2X7Rs was evident by a punctate staining pattern (Figure 3 C-D). Notably, the β-III tubulin antibody produced a more diffuse cellular staining compared to the noticeable punctate staining in the neuronal soma and along the processes by the P2X7R-specific antibody. Likewise, to assess the expression of P2X7Rs on hiPSC-derived astrocytes, double immunostaining was performed on astrocytes at 4 weeks *in vitro* using antibodies against the astrocyte marker GFAP and P2X7R. The punctate staining pattern for P2X7R was also present in GFAP-positive cells demonstrating the expression of P2X7Rs in astrocytes (Figure 3 E-H).

**Figure 3.**
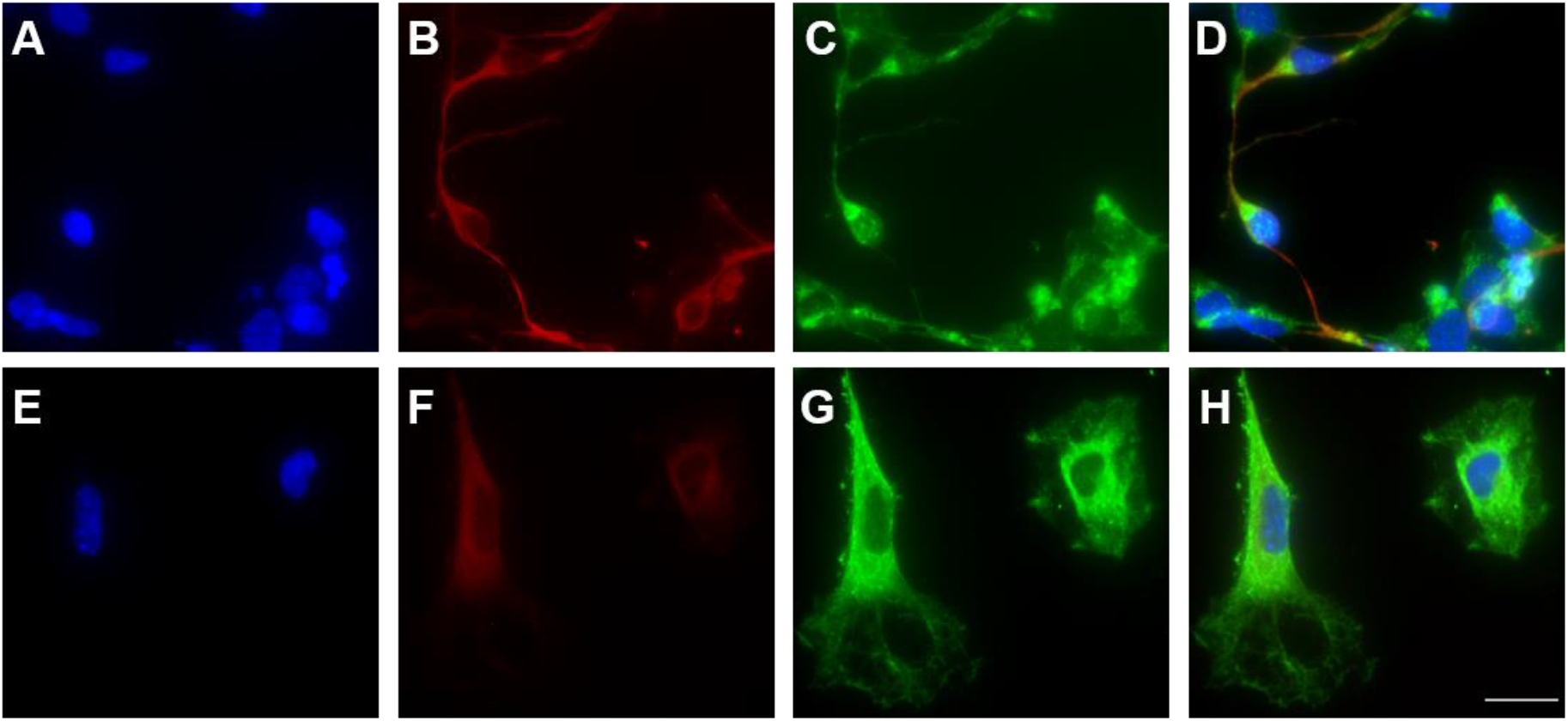
Immunofluorescence analysis of P2X7R expression in hiPSC-derived neurons and astrocyes. (**A–D**) Immunofluorescence analysis of neurons stained with an antibody against P2X7R (green) with double-labelling using neuronal marker β-III tubulin (red). (**E–H**) Immunofluorescence analysis of astrocyte marker GFAP. The nucleus is counterstained with Hoechst (blue). Note the punctate co-localization of P2X7R immunoreactivity with β -III tubulin or GFAP. Scale bar 50 µM.

These results demonstrate the expression of P2X7Rs on hiPSC-derived neuronal soma and processes as well as on astrocytes.

### 3.3. BzATP evokes AFC-5128-sensitive Ca^2+^ transients in hiPSC-derived neurons

To confirm the results obtained using immunocytochemistry, we next sought to assess whether P2X7Rs also respond to the application of P2X7R-stimulating agonists. Normally, the activation of the P2X7R requires high (in mM range) concentrations of ATP. The P2X7R is, however, 10 to 30 times more sensitive to the ATP analog and unselective P2X7R agonist, 2′,3′-O-(4-benzoylbenzoyl)-ATP (BzATP) [32, 33]. To assess the functional expression of P2X7Rs, hiPSC-derived neurons grown on coverslips were loaded with 2 μM Cal-520 AM calcium-sensitive fluorescent dye. Unstimulated neurons bathed in extracellular Cal-520 solution showed a measurable level of Cal-520 fluorescence (excitation 494 nm: emission 525 nm), indicating resting [Ca^2+^]i (Figure 3A).

The duration and time-course of the drug application was calibrated using fluorescein dye that was programmed to start at 5 s after the ejection of the control solution and continue to flow for a duration of 5 s. Analysis of the time-course of the fluorescein dye application confimed that the dye followed the preset stipulated time-couse (Figure 3B). During two consecutive appications of 300 μM BzATP, hiPSc-derived neurons responded to each application by a discernable increase in fluorescence (Figure 3C). To assess if the BzATP-evoked responses are P2X7R mediated, we repeated the BzATP application in the presence of the P2X7R antagonist AFC-5128 [34]. Fluorescence rose quickly in response to a pulse of 300 μM BzATP for 5 s signifying a persistent rise in [Ca^2+^]i during constant agonist exposure. In contrast, when the cells were pre-incubated with the P2X7R antagonist AFC-5128 (30 nM) pulse ejection of 300 μM BzATP and 30 nM AFC-5128 the change in [Ca^2+^]i was significantly reduced (Figure 3D). Individual neurons exhibited a synchronous increase in Cal-520 fluresence (ΔF) upon application of the agonist BzATP (Figure 3E) which was markedly reduced in the presence of the P2X7R inhibitior AFC-5128 (Figure 3F).

Taken together, these results show that the BzATP-gated change in [Ca^2+^]i in hiPSC-derived neurons is mediated via the P2X7R.

### 3.4. BzATP evokes AFC-5128-sensitive Ca^2+^ transients in iPSC-derived astrocytes

Similar to neurons, we then assessed the functional expression of P2X7Rs in hiPSC-derived astrocytes by monitoring changes in [Ca^2+^]i upon BzATP application, using the calcium-sensitive fluorescent reporter Cal-520. Unstimulated astrocytes bathed in extracellular solution showed a dim but measurable level of Cal-520 fluorescence (excitation 494 nm: emission 525 nm), indicating low resting [Ca^2+^]i (Figure 4A). The hiPSC-derived astrocytes exhibited spontaneous asynchronous [Ca^2+^]i fluctuations (Figure 4B). Similar to our findings in hiPSC-derived neurons, pulse ejection of BzATP (300 μM) for 5 s evoked an increase in [Ca^2+^]i in a population of hiPSC-derived astrocytes (Figure 4C).

**Figure 4.**
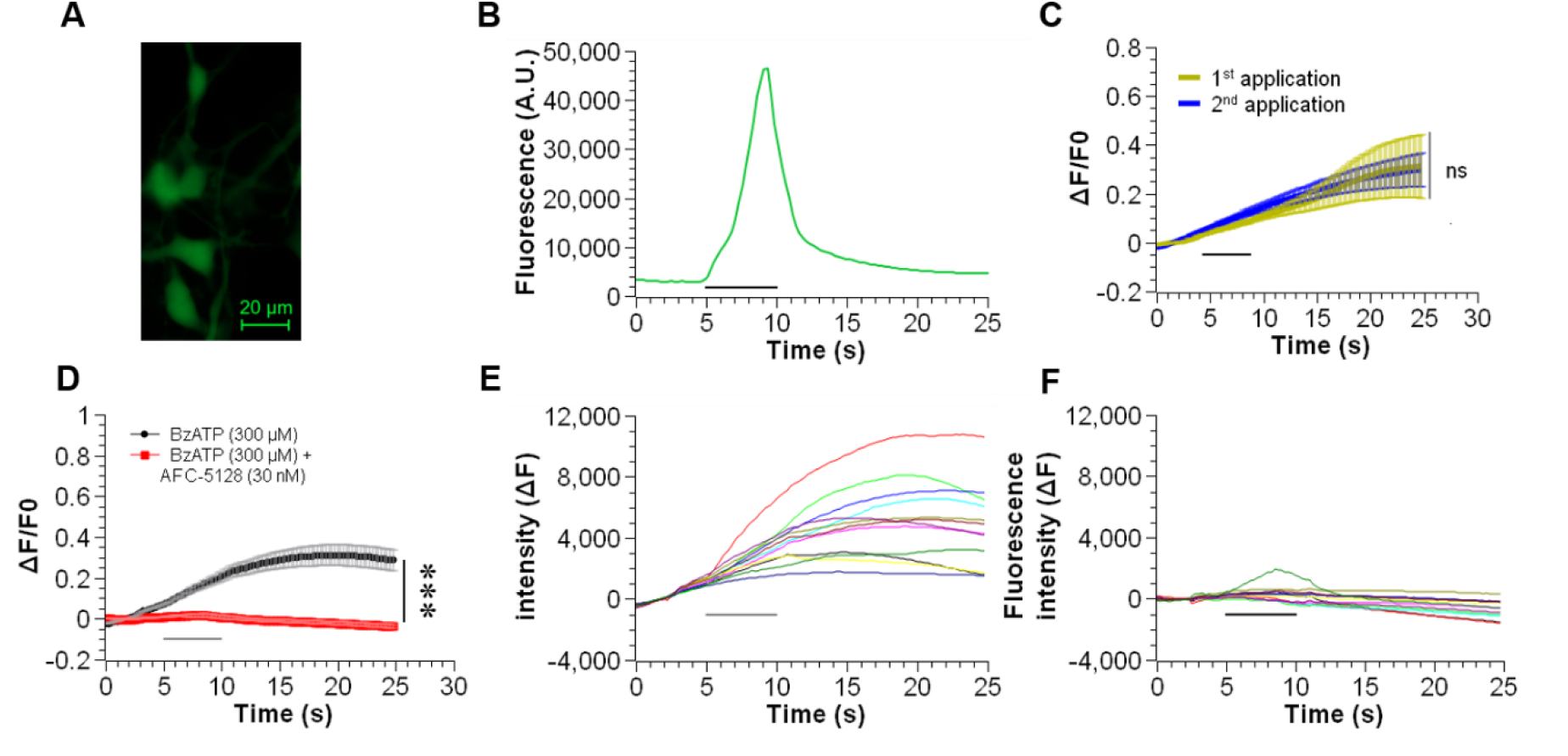
BzATP-evoked Ca^2+^ responses in hiPSC-derived neurons. (**A**) Representative image of Cal-520-loaded hiPSC-derived neurons at rest. (**B**) Graph illustrating the time course of fluorescein dye calibration. (**C**) BzATP-evoked calcium responses to two consecutive applications of 300 μM BzATP (Paired t-test, p> 0.05 (n = 8). (**D**) Response to the application of BzATP (300 μM) in the abcence (black trace) and presence (red trace) of the P2X7R antagonist AFC-5128 (30 nM). The values represent the mean and SEM. (Paired t-test, ***p < 0.001 (n = 14). (**E**) Traces illustrating BzATP -evoked calcium fluorescence response of individual neurons in the abcence of AFC-5128. (**F**) Traces illustrating BzATP -evoked calcium fluorescence response of individual neurons in the presence of AFC-5128.

**Figure 5.**
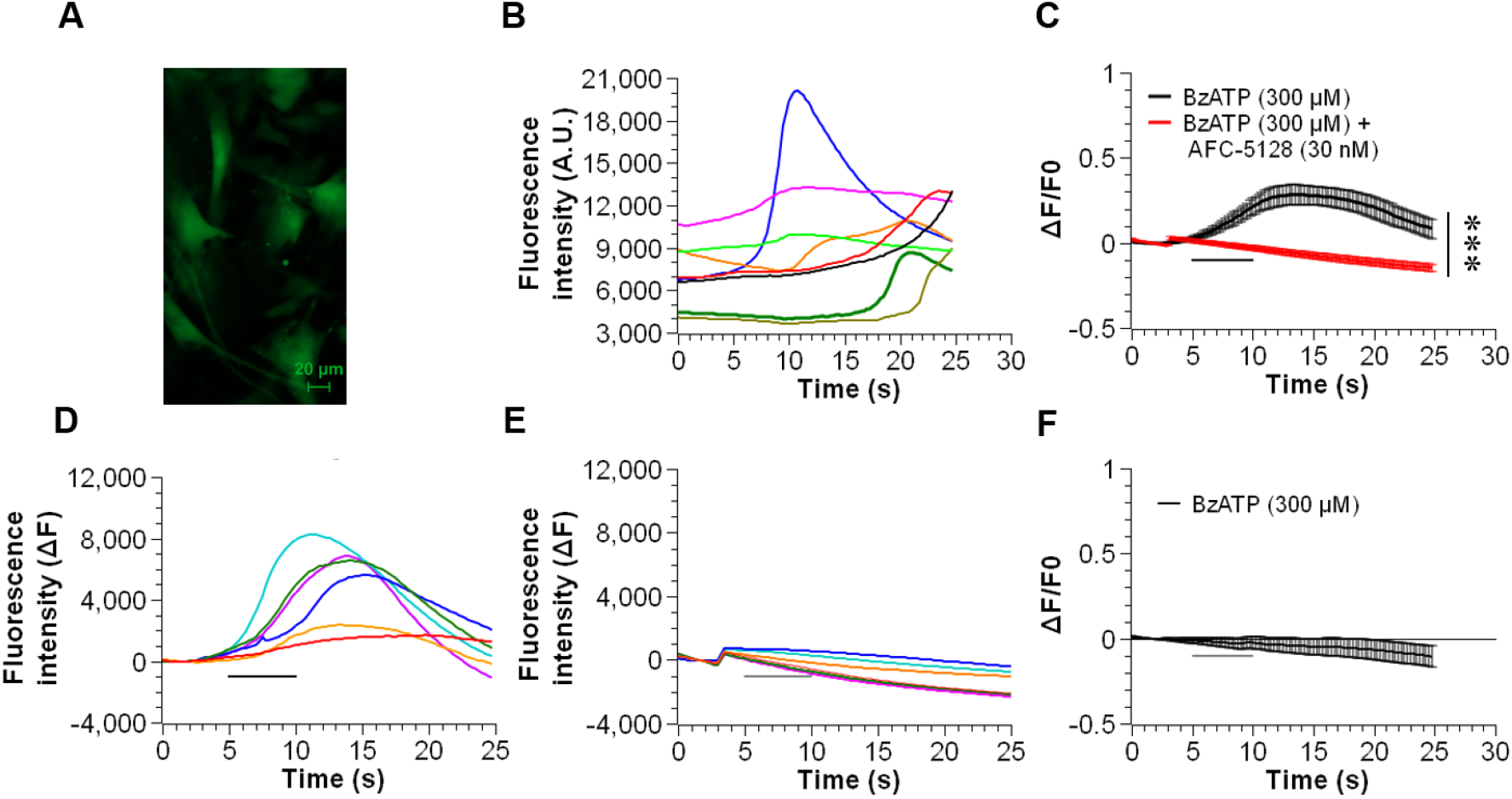
BzATP-evoked Ca2+ responses in hiPSC-derived neurons. (**A**) Representative image of Cal-520-loaded hiPSC-derived astrocytes at rest. (**B**) Exemplary traces depicting spontaneous activity in hiPSC-derived astrocytes. (**C**) Response to the application of BzATP (300 μM) in the abscence (black trace) and presence (red trace) of AFC-5128 (30 nM). The values represent the mean and SEM. (Paired t-test, ***p < 0.001 (n = 18). (**D**) Exemplary traces illustrating BzATP-evoked Cal-520 fluorescence responses of individual astrocytes in the abcence of AFC-5128 (30 nM). (**E**) Exemplary traces illustrating BzATP-evoked Cal-520 fluorescence responses of individual astrocytes in the presence of AFC-5128 (30 nM). (**F**) Response to the application of BzATP in the presence of 1 mM Mg^2+^ and 2 mM Ca^2+^.

As is typical of BzATP stimulation, this elevation in [Ca^2+^]i persisted for a few seconds after the agonist was washed out. In support of the role for the P2X7R in mediating this response, 3 min pre-incubation of astrocytes with the P2X7R antagonist AFC-5128, followed by co-application of BzATP with AFC-5128 (30 nM) significantly reduced BzATP-mediated [Ca^2+^]i response (Figure 4C). Individual astrocytes responded to BzATP in a synchronous manner by increasing their Cal-520 fluoresence (ΔF), indicative of increased [Ca^2+^]i (Figure 4D). This was markedly reduced in the presence of the P2X7R antagonist, AFC-5128 (Figure E). Physiological concentration of extracellular Mg^2+^ and Ca^2+^ is known to inhibit P2X7Rs [35]. Inhibition of the P2X7R by physiological concentrations of extracellular divalent cations is a characteristic feature of this receptor [36-38]. Consistent with these findings, we found that BzATP application in the presence of 1 mM Mg^2+^ and 2 mM Ca^2+^ failed to induce any decernable increase in Cal-520 fluorescence in hiPSC-derived astrocytes (Figure 4F). These results demonstrate that Ca^2+^ entry across the plasma membrane mediated via P2X7Rs contribute to the BzATP-evoked Ca^2+^ signals and confirms the functional expression of P2X7Rs in hiPSC-derived astrocytes.

## 4. Discussion

Here, we differentiated hiPSC-derived cortical forebrain neurons and glia exhibiting fundamental electrophysiological properties and calcium signaling which may make them a more suitable *in vitro* model for drug screening compared to current rodent models. Using fast agonist application and simultaneous Ca^2+^ imaging, we show that while both cell types, neurons and astrocytes, respond by [Ca^2+^]i increase upon BzATP stimulation, the selective P2X7R antagonist AFC-5128 abolishes this effect confirming the functional expression of P2X7R in hiPSC-derived neurons and astrocytes.

Despite extensive investigation into neuronal and glial P2X7R expression and function in both *in vitro* and *in vivo* animal models [39-41], limited studies have been conducted to characterize its expression and function in human-derived neurons and astrocytes and only two studies have reported the presence of *P2rx7* mRNA in cultured human adult and fetal brain tissue [29, 30]. P2X7R has been proposed as a potential therapeutic target in a plethora of human conditions ranging from cancer, to cardiovascular conditions to diseases of the CNS such as neurodegenerative diseases (e.g. Alzheimer’s disease), psychiatric diseases (e.g. schizophrenia, depression) and, more recently epilepsy [42-46]. Studies investigating a causative role of P2X7R during diseases have, however, been mainly restricted to experimental animal models. hiPSC-derived neurons and glia have been widely accepted as human brain-relevant *in vitro* cellular models to investigate phenotypic alterations in neurological diseases [25, 27, 47]. Another important consideration is the fact that P2X7R exhibits remarkable species-specific differences in their pharmacological properties to drugs such as AFC-5128. Thus, to advance findings from pre-clinical experimentation into the clinical setting, the development of human-derived cellular models for targeted drug screening, such as that for the P2X7R in this study, would serve an invaluable tool to bridge the gap between translational animal models and human neurological diseases.

Whereas the presence of P2X7R on microglia and oligodendrocytes is well-established, whether P2X7R is functional on neurons and astrocytes remains controversial [48, 49]. Data has suggested that on neurons, the main site for P2X7R seems to be at presynaptic terminals [50], where it may contributes to the regulation of neurotransmitter release, including GABA and glutamate [5]. Further evidence of neuronal expression of the P2X7R stems from studies using *P2rx7*-EGFP (enhanced green fluorescent protein) reporter mice [51] and mice expressing humanized P2X7R [52]. Other studies have, however, failed to detect P2X7R on neurons [53, 54]. P2X7R has also been shown to be expressed and functional on astrocytes *in vitro* and *in vivo [41]*. Moreover, P2X7R expression has been shown to be increased in several disease models including in ischemia and transgenic models of Alzheimer’s disease [55, 56]. Similar to neurons, conflicting evidence exists on whether P2X7Rs are expressed on astrocytes [54]. Our findings that both hiPSC-derived neurons and astrocytes show surface expression of P2X7Rs has wide implications in signaling mechanisms in the brain. Both neurons and astrocytes release ATP which is the endogenous ligand for P2X7Rs [57]. Although ATP is a low affinity ligand at P2X7Rs, [57] P2X7Rs on astrocytes may be activated by ATP released from both neurons and astrocytes leading to signal amplification. P2X7R activation on astrocytes could promote enhanced astrocyte-mediated glutamate release leading to increased network excitability. Likewise, ATP released from astrocytes could also directly activate neuronal P2X7Rs promoting hyper-excitable neuronal networks. In addition, at the site of brain injury, exogenous ATP mediates the release of pro-inflammatory cytokines leading to neuroinflammation via P2X7Rs [58] possibly via upregulation of neuronal and/or astrocytic P2X7R during neurological conditions.

While our results suggest P2X7Rs to be functional on both human-derived astrocytes and neurons, it is important to keep in mind that our studies are carried out under *in vitro* conditions and whether P2X7Rs are expressed *in vivo* in humans requires further investigation and clarification including, for example patch clamp in human brain slices. Our results do not show what neuronal subtype expresses P2X7Rs, although their activation is known to affect both glutamate and GABA neurotransmission, suggesting their expression on excitatory and inhibitory neurons [5]. Differentiation of layer-specific cortical neurons from hiPSCs has been well established [18, 59, 60], allowing elucidation of the functional expression of P2X7Rs in layer-specific cortical GABAergic and glutamatergic neurons derived from hiPSCs in future studies. P2X7Rs have been shown to be upregulated in human reactive astrocytes [30]. We did not detect discernable changes in morphology of astrocytes in culture over time, but did not assess if the culture conditions leads to neuroinflammation and the generation of reactive astrocytes.

Finally, BzATP-evoked responses should be interpreted with caution because BzATP is a non-specific agonist at purinergic receptors and agonist activity at non-P2X7Rs has also been documented. Examples include BzATP-evoked responses at ionotropic P2X1, P2X2 and P2X3 [61] and metabotropic P2Y2, P2Y11 and P2Y13 [62, 63]. In addition, BzATP is metabolized to other adenine derivatives like Bz-adenosine by ecto-nucleotidases and could activate A1 receptors in tissues [64]. Action by BzATP metabolites is, however, highly unlikely because the drug is superfused onto the recorded cells as a continuous stream of fast flowing solution and quickly removed from the imaging chamber. Importantly, suggesting BzATP-mediated responses are mainly mediated mainly via P2X7Rs, AFC-5128, specific P2X7R antagonist [34], suppressed calcium increases both in neurons and astrocytes.

In conclusion, here we demonstrate the expression of functional P2X7Rs on hiPSCs-derived neurons and astrocytes contributing further evidence for the expression of P2X7Rs in these cell types and providing the proof of concept that hiPSCs represent a valid model to study P2X7Rs signaling in a human cellular model.

## Author Contributions

Conceptualization, J.K. and T.E.; methodology, J.K., T.E.; formal analysis, J.K.; investigation, J.K.; resources, T.E., O.W., K.D., M.H., D.H., J.H.M.P.; writing—original draft preparation, J.K., T.E.; writing—review and editing, J.K., O.W., T.E., D.H., J.H.M.P.; supervision, T.E.; project administration, T.E.; funding acquisition, J.K., T.E., D.H. and J.H.M.P. All authors have read and agreed to the published version of the manuscript.

## Acknowledgments

The authors acknowledge Dr. Gareth Morris for the support during the establishment of the iPSC facility.

